# Left Temporoparietal Neural Synchrony Tracks Cooperative Performance Across Adult Dyads

**DOI:** 10.64898/2026.10.20.760341

**Authors:** Alessandro Carollo, Riccardo Cavallaro, Andrea Bizzego, Dagmara Dimitriou, Stefanie Hoehl, Gianluca Esposito

**Affiliations:** Department of Psychology and Cognitive Science, University of Trento, 38068 Rovereto, Italy; Sleep Education and Research Laboratory, Department of Psychology and Human Development, UCL Institute of Education, University College London, London, WC1H 0AA, UK; Department of Developmental and Educational Psychology, University of Vienna, 1010 Vienna, Austria

## Abstract

Cooperation requires interacting individuals to coordinate their actions toward shared goals, and interpersonal neural synchrony (INS) has been proposed as a neural correlate of this coordination. However, the relationship between INS and cooperative performance remains heterogeneous, and relatively little is known about how it varies across social relationships. The present study examined INS and behavioral performance during a naturalistic cooperative tabletop Jenga task in close-friend, romantic-partner, and mother–child dyads (N = 284 participants). Brain activity was simultaneously recorded using functional near-infrared spectroscopy (fNIRS) hyperscanning, and INS was quantified using wavelet transform coherence across the bilateral inferior frontal gyrus (IFG) and temporoparietal junction (TPJ). Cooperative performance was assessed using tower collapse and the number of valid turns completed during the game. True interacting dyads exhibited significantly greater neural synchrony than surrogate dyads across all examined regional pairings (***d_range_*** = 0.35–0.55), indicating robust interaction-related inter-brain synchrony. Behavioral performance was not associated with INS at the whole-network level. Region-specific analyses, however, showed that tower collapse was associated with reduced synchrony in the left TPJ–left TPJ (***p*_Bonf_** = 0.009) and left TPJ–right TPJ (***p*_Bonf_** = 0.017) pairings, whereas the number of valid turns was unrelated to INS. Follow-up analyses further indicated that the association between tower collapse and left TPJ–left TPJ synchrony was present in close-friend (***p*_Bonf_** = 0.020) and romantic-partner (***p*_Bonf_** = 0.026) dyads but not in mother–child dyads. These findings suggest that the behavioral relevance of interpersonal neural synchrony is not uniform, but depends on the neural systems involved, the specific index of cooperative performance, and the relational and developmental context in which interaction unfolds.

---

Keywords: cooperation, cooperative performance, fNIRS, hyperscanning, interpersonal neural synchrony, temporoparietal junction

## 1 Introduction

Cooperation is a fundamental organizing principle of human action (Nowak, 2006; Rand & Nowak, 2013). Achieving shared goals requires individuals to contribute their own skills and effort and to continuously coordinate their actions with those of others. Cooperative performance may therefore emerge, at least in part, from the ability of interacting partners to monitor, predict, and adapt to one another during ongoing joint action (Shamay-Tsoory, Saporta, Marton-Alper, & Gvirts, 2019). Successful coordination also requires partners to flexibly adapt their actions to one another, and such adaptation can signal a willingness to invest effort in the interaction, thereby strengthening commitment to the joint activity (McEllin, Felber, & Michael, 2023).

One prominent manifestation of interpersonal coordination is synchrony, broadly referring to the temporal alignment of events between interacting individuals (Hamilton et al., 2026; Hoehl, Fairhurst, & Schirmer, 2021). Behavioral synchrony has been linked to important cooperative outcomes. Experimentally inducing synchronous action has been shown to increase subsequent cooperation and prosocial behavior (Kirschner & Tomasello, 2010; Marsh, Richardson, & Schmidt, 2009; Wiltermuth & Heath, 2009), and may also enhance individuals’ sensitivity to their partner’s actions, thereby facilitating performance in joint-action tasks (Valdesolo, Ouyang, & DeSteno, 2010). Thus, interpersonal synchrony may both emerge from and contribute to the processes that allow interacting individuals to coordinate effectively toward a shared goal.

Interpersonal synchrony extends beyond overt behavior (Carollo, 2026; Feldman, 2014, 2017). During social interactions, coordinated dynamics have also been observed at the physiological level, including the alignment of autonomic and peripheral signals across interacting partners (Gordon & Bartsch, 2026). Similar forms of interpersonal coordination can also emerge at the neural level, with the activity of interacting individuals becoming dynamically aligned over the course of an interaction (De Felice et al., 2025). This phenomenon, commonly referred to as interpersonal neural synchrony, has been increasingly investigated using hyperscanning approaches that record brain activity simultaneously from two or more interacting individuals (Carollo & Esposito, 2024; Czeszumski et al., 2020; Dumas, Nadel, Soussignan, Martinerie, & Garnero, 2010; Montague et al., 2002). Interpersonal neural synchrony has been observed across a range of social contexts, including communication, joint attention, imitation, and cooperation, suggesting that inter-brain coupling may reflect aspects of the dynamic interpersonal coordination involved in successful joint action (De Felice et al., 2025).

Beyond its association with cooperation, interpersonal neural synchrony has also been linked to how effectively individuals perform together (Gordon & Bartsch, 2026). Across a variety of joint-action paradigms, greater inter-brain alignment has been associated with more successful behavioral coordination and better joint performance (Czeszumski et al., 2020; Hoehl et al., 2021). A meta-analysis of hyperscanning studies found an overall positive association between interpersonal neural synchrony and team performance across a range of collaborative contexts (Ŕeveilĺe, Vergotte, Perrey, & Bosselut, 2024). Consistent with this pattern, early fNIRS hyperscanning work showed that interpersonal synchrony in the right superior frontal cortex increased specifically during cooperation and was associated with better cooperative performance (Cui, Bryant, & Reiss, 2012). Similarly, during a cooperative tangram task, mother–child dyads exhibited greater neural synchrony in prefrontal and temporo-parietal regions when solving problems together than individually, and greater synchrony predicted joint problem-solving success above and beyond behavioral reciprocity (Nguyen et al., 2020). Together, these findings suggest that interpersonal neural synchrony is closely associated with the effectiveness of ongoing joint coordination.

However, the relationship between interpersonal neural synchrony and performance is far from uniform (Gordon & Bartsch, 2026). Hyperscanning studies have employed highly heterogeneous interactive paradigms, ranging from economic and turn-taking games to joint problem solving, musical interaction, and naturalistic games such as Jenga, and the neural patterns observed appear to depend considerably on the specific demands and structure of the interaction (Carollo, Bizzego, et al., 2025; Czeszumski et al., 2020). Importantly, such variability may also reflect differences in who is interacting (Gordon & Bartsch, 2026). Dyads differ in familiarity, interpersonal closeness, shared history, previous experience coordinating with one another, and the social meaning attributed to the interaction. These relational characteristics may shape how partners anticipate, interpret, and adapt to one another’s actions, thereby influencing both joint performance and the neural dynamics accompanying coordination. Consistent with this possibility, interpersonal neural synchrony and cooperative behavior have been shown to vary across relationship types, with romantic partners, friends, and unfamiliar partners exhibiting distinct patterns during comparable cooperative interactions (Carollo, Bizzego, et al., 2025; Djalovski, Dumas, Kinreich, & Feldman, 2021; Pan, Cheng, Zhang, Li, & Hu, 2017). This heterogeneity is also reflected in theoretical accounts: whereas earlier perspectives emphasized synchrony as an adaptive feature of social affiliation and coordination (Feldman, 2014, 2017), more recent frameworks emphasize that its functional significance may depend on the interactional and relational context (De Felice et al., 2025; Gordon, Tomashin, & Mayo, 2025).

Despite this growing emphasis on context, the role of relational relationships in shaping the association between interpersonal neural synchrony and cooperative performance remains poorly understood. Cooperative behavior does not occur in a relational vacuum, as the same joint task may involve different expectations, coordination strategies, and interpersonal significance depending on the relationship between the individuals involved. For instance, close friends, romantic partners, and parent– child dyads are characterized by distinct relational histories and interaction dynamics, which may affect both how effectively partners coordinate and how such coordination is reflected at the neural level. Understanding interpersonal neural synchrony in joint action may therefore require considering not only whether synchrony is associated with successful performance, but also how this association is expressed across different relational contexts.

The present study addresses this question in a naturalistic cooperative setting by examining close friends, romantic partners, and mother–child dyads while they perform a cooperative tabletop Jenga task, included in the study by Carollo, Bizzego, et al. (2025). First, we hypothesized that true interacting dyads would exhibit greater interpersonal neural synchrony than surrogate dyads, reflecting a co-presence effect on brain synchrony. Second, based on accounts emphasizing the adaptive role of interpersonal synchrony, we hypothesized that greater neural synchrony would be associated with better cooperative performance, reflected in a lower likelihood of tower collapse and a greater number of valid turns. Finally, motivated by more recent context-dependent accounts of interpersonal synchrony (De Felice et al., 2025; Gordon et al., 2025), we examined whether the pattern of the synchrony–performance association differed across close-friend, romantic-partner, and mother–child dyads. By comparing the synchrony–performance relationship within multiple forms of close relationships using the same interactive paradigm, the study aims to provide insight into how relational context may contribute to the neural dynamics associated with successful real-time cooperation.

## 2 Methods

### 2.1 Study design

Data were collected as part of a broader hyperscanning study in which participant dyads completed multiple social interaction tasks while their brain activity was simultaneously recorded using fNIRS (Carollo, Bizzego, et al., 2025). The project included three dyad types: close friends, romantic partners, and mother-child pairs, selected to represent distinct forms of close social relationships that are frequently investigated in the interpersonal synchrony and developmental literature. Data from close friends and romantic partners were collected at the University of Trento (Italy), whereas data from mother-child dyads were collected at the University of Vienna (Austria).

During the experimental session, dyads completed three social tasks differing in their degree and structure of interpersonal interaction: video co-exposure, a rule-based cooperative game, and free conversation. The present study focuses exclusively on the rule-based cooperative condition, consisting of a 5-min cooperative Jenga task, to examine the association between interpersonal neural synchrony and cooperative performance across different dyadic relationships.

The study received ethical approval from the Ethics Committees of the University of Trento (ref. 2022-052) and the University of Vienna (ref. 00732) and was conducted in accordance with the Declaration of Helsinki. Written informed consent was obtained from all adult participants. For child participants, informed consent was provided by their parent or legal guardian.

### 2.2 Participants

A total of 284 participants, forming 142 dyads, took part in the study. The sample comprised 70 close-friend dyads (29 male–male, 23 female–female, and 18 mixed-gender dyads; *M*_age_ = 21.60 *±* 2.06 years), 39 heterosexual romantic-partner dyads (*M*_age_ = 23.45 *±* 3.08 years), and 33 mother–child dyads (20 female children; mothers: *M*_age_ = 38.59 *±* 4.66 years; children: *M*_age_ = 5 years, 6 months, 18 days *±* 3 months, 22 days). Two close-friend dyads, one romantic-partner dyad, and two mother–child dyads were excluded because of technical issues during data collection.

Close-friend dyads were required to consist of individuals who identified as close friends and had no current or previous romantic relationship with one another. Romantic-partner dyads consisted of heterosexual couples who were in an ongoing romantic relationship at the time of data collection. Mother-child dyads included mothers with children aged 5 years. Across all groups, participants reported no known or diagnosed medical or neurological conditions, particularly conditions that could affect cerebral oxygenation or the oxygen-binding capacity of the blood.

Close-friend and romantic-partner dyads were recruited through convenience and snowball sampling, primarily via social media and the University of Trento participant recruitment system (SONA). Mother-child dyads were recruited from an existing volunteer database at the University of Vienna. Close-friend and romantic-partner dyads received no financial compensation. Mother-child dyads received €6 to cover transportation costs and a small toy for the child (approximately €6 in value).

### 2.3 Experimental tasks

All dyads began the experimental session with a resting-state baseline recording (2 min for adult dyads and 1 min for mother–child dyads). Mother–child dyads additionally completed a 1-min resting-state recording between experimental tasks. Participants were seated in separate chairs approximately 60–80 cm apart, with the same general configuration maintained across dyad types and experimental conditions.

As part of the experimental protocol, dyads completed three social tasks: video co-exposure, a rule-based cooperative game, and free verbal interaction. The order of the three tasks was randomized across dyads to minimize potential order effects. The present study focuses exclusively on the rule-based cooperative condition, consisting of a 5-min cooperative Jenga task, and analyses were conducted on the full 5-min interaction.

During the Jenga task, a tower was positioned on a table between the two participants, who sat facing each other while their brain activity was simultaneously recorded using fNIRS (Figure 1A). Dyad members were instructed to take turns removing one block at a time and placing it on top of the tower. To maintain comparable task difficulty across dyads, participants were instructed to remove only one block from each layer, starting from the bottom of the tower. If the tower collapsed before the end of the task, participants reconstructed it and resumed playing so that the total interaction duration remained fixed at 5 min for all dyads. The task therefore provided a semi-structured cooperative interaction in which participants worked toward a shared goal within a common set of rules and a predictable turn-taking structure.

**Fig. 1.**
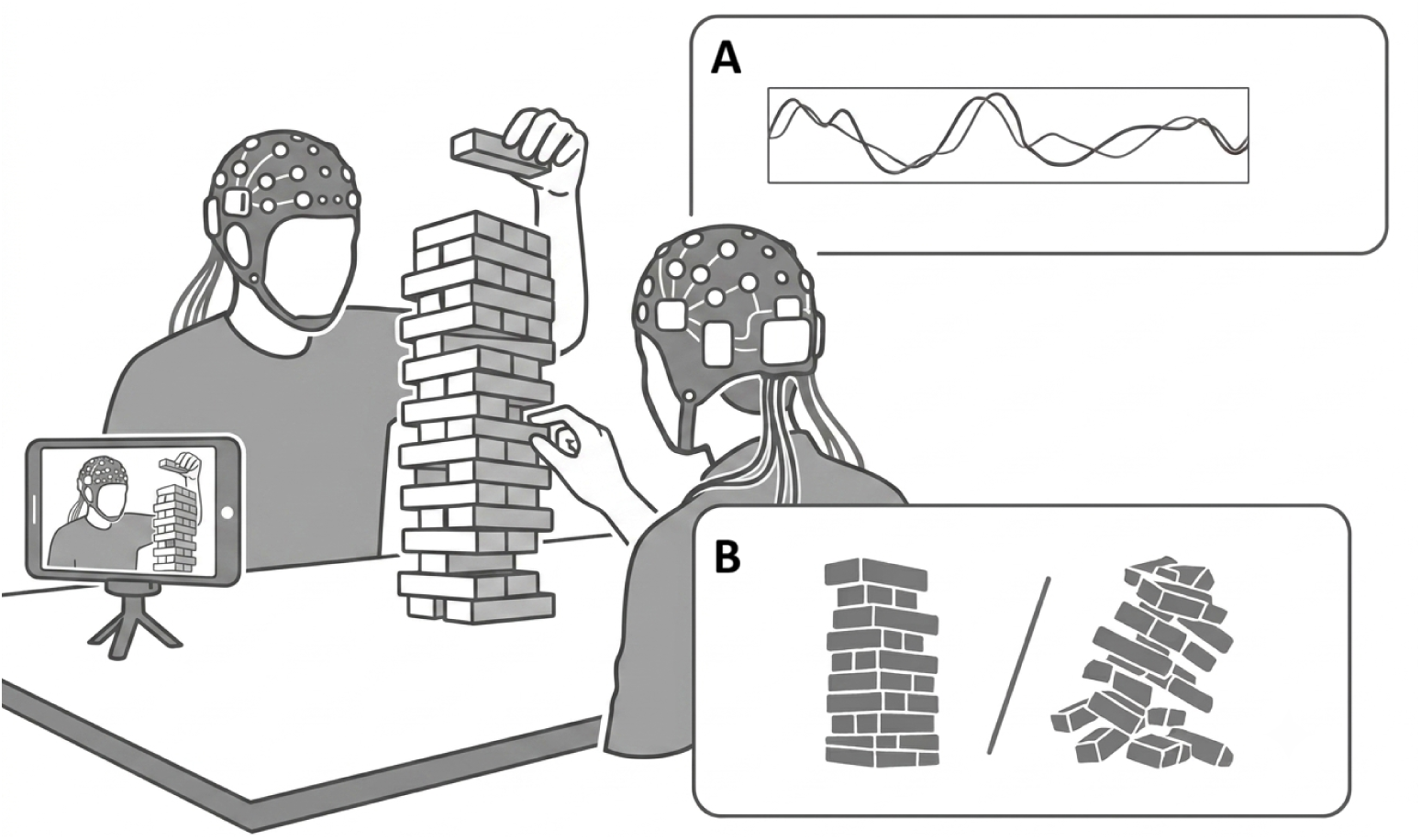
Experimental setup and behavioral coding during the cooperative Jenga task. **(A)** Participants sat facing each other while their brain activity was simultaneously recorded using fNIRS. **(B)** The interaction was also video-recorded to enable behavioral coding of task performance, including events such as tower collapse.

Behavioral performance during the Jenga interaction was subsequently coded from the task recordings based on game outcomes, including the number of successfully completed levels and tower collapses. The behavioral coding procedure and derivation of the performance measures are described in detail below.

### 2.4 Behavioral coding

The cooperative Jenga task was video-recorded to enable behavioral coding of task performance (Figure 1B), while participants’ brain activity was simultaneously recorded using fNIRS. Behavioral performance was coded separately for each dyad from the video recordings.

Task performance was quantified as the total number of valid turns completed during the 5-min interaction. A turn was considered valid when a participant successfully removed one wooden block from the tower and placed it on top without causing the tower to collapse. The total number of valid turns therefore indexed the number of successfully completed game actions within the fixed task duration, with higher values indicating better cooperative task performance. This criterion was applied consistently across dyads to exclude incomplete, irregular, or otherwise ambiguous actions from the performance measure.

Tower collapse was coded separately as a binary outcome indicating whether the tower collapsed at least once during the interaction (*fallen* vs. *not fallen*). Following a collapse, participants rebuilt the tower and resumed the game until the end of the 5-min task, allowing performance to be assessed over the full interaction period.

### 2.5 Acquisition of neural data

Neural activity was recorded using an fNIRS hyperscanning setup. Each participant wore an optode cap equipped with eight light sources emitting at wavelengths of 760 and 850 nm and eight detectors. The optode configuration was designed to measure hemodynamic activity over the bilateral inferior frontal gyrus (IFG) and temporoparietal junction (TPJ), consistent with previous hyperscanning studies (e.g., Lei, Liu, Guo, Zuo, & Wen, 2025; Nguyen et al., 2020).

Optode configurations differed slightly between data collection sites. For close-friend and romantic-partner dyads recorded in Italy, one detector was dedicated to short-separation measurements, yielding eight short-distance channels and 14 long-distance fNIRS channels per participant. For mother–child dyads recorded in Vienna, short-separation channels were not available; instead, the additional detector was used to acquire two further long-distance channels, resulting in 16 channels per participant. For subsequent analyses, long-distance fNIRS channels were grouped into four regions of interest (ROIs): left IFG, right IFG, left TPJ, and right TPJ. For close-friend and romantic-partner dyads, the optode layout was optimized using the fNIRS Optodes’ Location Decider (fOLD) to maximize sensitivity to the targeted cortical regions (Zimeo Morais, Balardin, & Sato, 2018). For mother–child dyads, the cap configuration followed previously established adult–child hyperscanning protocols (Nguyen et al., 2020; Nguyen, Schleihauf, Kayhan, et al., 2021; Nguyen, Schleihauf, Kungl, et al., 2021).

Optode placement was guided by the international 10–20 EEG system. During data collection in Italy, optode stabilizers were used to maintain source–detector distances of no more than 3 cm. Data acquisition systems also differed between sites: close-friend and romantic-partner dyads were recorded using a NIRSport2 system (NIRx Medical Technologies LLC) at a sampling rate of 10.17 Hz, whereas mother–child dyads were recorded using a NIRSport system (NIRx Medical Technologies LLC) at 7.81 Hz.

### 2.6 Processing of neural data

Preprocessing and synchrony analyses were performed using *pyphysio* (Bizzego, Battisti, Gabrieli, Esposito, & Furlanello, 2019). For the present study, preprocessing was conducted on the full 5-min recording of the cooperative Jenga task. Raw fNIRS intensity signals were first converted into changes in optical density. Signal quality was then assessed using a convolutional neural network trained to classify the quality of fNIRS signal segments (Bizzego, Gabrieli, & Esposito, 2021; Bizzego, Neoh, Gabrieli, & Esposito, 2022).

The resulting signals were resampled to 10 Hz and converted into concentration changes in oxygenated (HbO) and deoxygenated hemoglobin (HbR) using the modified Beer–Lambert law (Delpy et al., 1988). Consistent with previous hyperscanning research, the analyses reported in the main text focused on HbO signals (Lim, Carollo, Bizzego, Chen, & Esposito, 2024a, 2024b), whereas corresponding analyses based on HbR signals are reported in the Supplementary Materials.

Finally, the hemoglobin signals were pre-whitened using an autoregressive model to reduce temporal autocorrelation and attenuate physiological noise (Barker, Aarabi, & Huppert, 2013). Pre-whitening prior to coherence analysis has been shown to reduce inflated coherence estimates and the associated risk of false-positive inter-brain coupling (Santosa, Aarabi, Perlman, & Huppert, 2017). Periods following tower collapse, during which participants reconstructed the tower and resumed the game, were retained as part of the 5-min interaction and were therefore included in the neural synchrony analyses.

### 2.7 Interpersonal neural synchrony

Interpersonal neural synchrony was estimated using wavelet transform coherence (WTC) (Carollo, Soffner, Esposito, & Bizzego, 2026; Grinsted, Moore, & Jevrejeva, 2004). WTC quantifies the coupling between two time series jointly in the time and frequency domains and can capture both in-phase and phase-lagged relationships (Nguyen et al., 2020).

For each dyad, WTC was computed between all possible pairs of good-quality long-distance fNIRS channels across the two participants. Thus, each good-quality channel from one dyad member was paired with each good-quality channel from the other dyad member, rather than restricting the analysis to spatially corresponding or homologous channels. Each resulting channel-pair observation was subsequently classified according to the regions of interest to which the two channels belonged: left IFG, right IFG, left TPJ, or right TPJ.

This procedure resulted in 10 unique unordered ROI pairings: four homologous pairings (left IFG–left IFG, right IFG–right IFG, left TPJ–left TPJ, and right TPJ– right TPJ) and six non-homologous pairings. For non-homologous combinations, the ordered configuration of the two ROIs was additionally retained to distinguish, for example, a left IFG channel from participant A paired with a left TPJ channel from participant B from the reverse configuration. Importantly, WTC estimates were retained at the individual channel-pair level for statistical analyses rather than being averaged into a single value for each ROI pairing.

For each channel pair, WTC was computed over the full 5-min Jenga interaction across a frequency range of 0.01–0.20 Hz, consistent with previous hyperscanning studies (Carollo, Stella, Lim, Bizzego, & Esposito, 2025; Lim et al., 2024a, 2024b). Coherence values were then averaged across this frequency range to obtain a single WTC estimate for each channel pair. This approach avoided restricting the analysis to a narrower frequency band in the absence of an *a priori* hypothesis regarding the frequency range in which synchrony would emerge (Carollo, Stella, et al., 2025).

To estimate a baseline level of inter-brain coherence in the absence of direct interaction, WTC was also computed for surrogate dyads. Surrogate dyads were generated by randomly pairing participants belonging to different original dyads within the same dyad type and data-collection cohort, thereby avoiding cross-group and cross-site pairings. Following Carollo, Stella, et al. (2025), a single permutation was performed, resulting in the same number of surrogate and true dyads.

For each surrogate dyad, WTC was computed using the same all-to-all channel-pairing procedure applied to true dyads. Each good-quality channel from one participant was paired with each good-quality channel from the participant drawn from another original dyad, and the resulting channel pairs were classified according to the same ROI-pairing scheme. Thus, surrogate participants completed the same task under comparable recording conditions but did not interact with one another. Synchrony estimated from these surrogate pairings provides a reference level of inter-brain coherence that can arise in the absence of real-time interpersonal interaction, including coupling related to common task structure and shared signal characteristics. Comparison with true dyads therefore allows interaction-related synchrony to be distinguished from this non-interactive baseline (Golland, Arzouan, & Levit-Binnun, 2015).

### 2.8 Data Analysis

Statistical analyses were conducted using linear mixed-effects models implemented in the *lme4* and *lmerTest* packages in R. Analyses proceeded in three stages.

First, we tested whether interpersonal neural synchrony during the Jenga task exceeded the level of inter-brain coherence observed in non-interacting surrogate dyads. Channel-pair WTC estimates were grouped into the 10 unique unordered ROI pairings. Separate mixed-effects models were fitted for each ROI pairing, with WTC at the channel-pair level as the dependent variable and dyad type (*true* vs. *surrogate*) as a fixed effect. For homologous ROI pairings, models included random intercepts for cohort and dyad. For non-homologous pairings, additional random intercepts accounted for the ordered ROI configuration and the assignment of the respective regions to dyad members. *p*-values for the true-versus-surrogate comparison were corrected across the 10 ROI-pair models using the Bonferroni procedure.

Second, we examined whether interpersonal neural synchrony was associated with behavioral performance during the Jenga task. These analyses were restricted to true dyads. The analysis was conducted at the channel-pair level: each WTC estimate obtained from a pair of channels constituted one observation, and channel-pair observations were not averaged within ROIs before model fitting. The number of valid turns completed during the task and tower collapse (*fallen* vs. *not fallen*) were entered simultaneously as fixed effects. Random intercepts were included for cohort, ordered ROI configuration, unordered ROI pairing, and dyad to account for the repeated channel-pair observations and their hierarchical organization within regional combinations and dyads. The same associations were subsequently examined separately within close-friend, romantic-partner, and mother–child dyads to characterize relationship-specific patterns.

Third, we examined whether associations between Jenga performance and interpersonal neural synchrony were specific to particular regional combinations. Channel-pair observations were divided according to the 10 unique unordered ROI pairings, and a separate mixed-effects model was fitted for each pairing. WTC at the channel-pair level was entered as the dependent variable, with number of valid turns and tower collapse as fixed effects. Models included random intercepts for cohort and dyad; for non-homologous ROI pairings, ordered ROI configuration was additionally included as a random effect to account for the two possible assignments of the regions across dyad members. *p*-values were corrected for multiple comparisons across ROI-level analyses using the Bonferroni procedure. ROI pairings showing evidence of an association with behavioral performance were subsequently examined separately within each relationship type to characterize cohort-specific effects.

## 3 Results

### 3.1 Behavioral performance across dyad types

Descriptive statistics for Jenga performance are reported in Table 3.1. The number of valid turns was broadly comparable across dyad types, with close-friend dyads completing an average of 11.5 valid turns (*SD* = 4.14), romantic-partner dyads 12.2 (*SD* = 4.22), and mother–child dyads 12.0 (*SD* = 4.91). Tower collapse occurred in 16.2% of close-friend dyads, 28.2% of romantic-partner dyads, and 66.7% of mother– child dyads.

**Table 1.** Descriptive statistics for cooperative Jenga performance across dyad types.

| Dyad type | Valid turns, $M$ ($SD$) | Median [IQR] | Tower collapse, $n$ (%) |
| --- | --- | --- | --- |
| Close friends | 11.5 (4.14) | 11 [7.00] | 11 (16.2) |
| Romantic partners | 12.2 (4.22) | 13 [6.00] | 11 (28.2) |
| Mother-child | 12.0 (4.91) | 11 [6.75] | 20 (66.7) |

**Table 2.** Comparison of interpersonal neural synchrony between true and surrogate dyads across ROI pairings during the cooperative Jenga task. Positive coefficients indicate greater wavelet transform coherence (WTC) in true compared with surrogate dyads. *P* values were Bonferroni-corrected across the 10 ROI comparisons. (*** *p*_Bonf_ *< .*001).

| ROI pairing | $\beta$ | SE | $t$ | df | $p_{\text{Bonf}}$ | $d$ [95% CI] |
| --- | --- | --- | --- | --- | --- | --- |
| Left IFG – Left IFG | 0.00395 | 0.00025 | 15.74 | 3922.3 | < .001*** | 0.50 [0.43, 0.56] |
| Left IFG – Right IFG | 0.00418 | 0.00019 | 21.53 | 7856.6 | < .001*** | 0.48 [0.44, 0.53] |
| Left IFG – Left TPJ | 0.00408 | 0.00019 | 21.29 | 7754.8 | < .001*** | 0.48 [0.44, 0.53] |
| Left IFG – Right TPJ | 0.00387 | 0.00024 | 16.01 | 4595.2 | < .001*** | 0.47 [0.41, 0.53] |
| Right IFG – Right IFG | 0.00471 | 0.00028 | 17.00 | 3806.5 | < .001*** | 0.55 [0.48, 0.61] |
| Right IFG – Left TPJ | 0.00419 | 0.00020 | 20.69 | 7630.1 | < .001*** | 0.47 [0.43, 0.52] |
| Right IFG – Right TPJ | 0.00410 | 0.00025 | 16.14 | 4501.0 | < .001*** | 0.48 [0.42, 0.54] |
| Left TPJ – Left TPJ | 0.00427 | 0.00027 | 15.70 | 3713.8 | < .001*** | 0.52 [0.45, 0.58] |
| Left TPJ – Right TPJ | 0.00299 | 0.00026 | 11.68 | 4441.2 | < .001*** | 0.35 [0.29, 0.41] |
| Right TPJ – Right TPJ | 0.00444 | 0.00043 | 10.27 | 1354.6 | < .001*** | 0.54 [0.32, 0.76] |

**Table 3.**
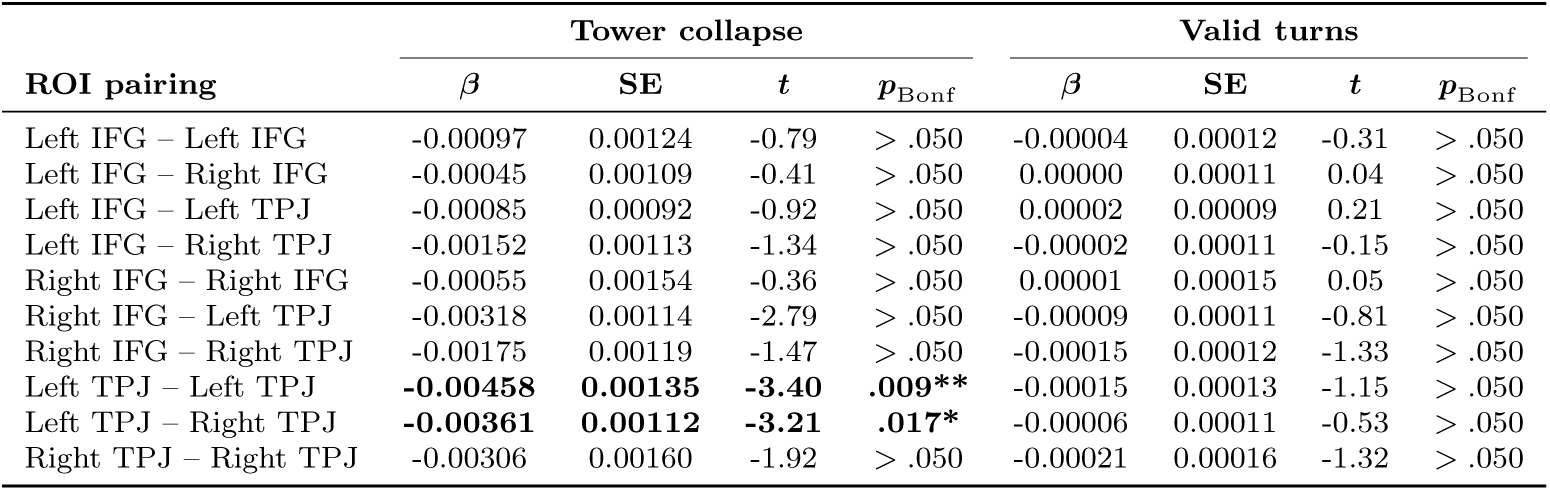
Associations between behavioral performance and interpersonal neural synchrony across ROI pairings. Tower collapse (*fallen* vs. *not fallen*) and the number of valid turns were entered simultaneously as predictors of wavelet transform coherence (WTC). *P* values were Bonferroni-corrected separately for each behavioral predictor across the 10 ROI pairings. Significant effects after correction are shown in bold (* *p*_Bonf_ *< .*050; ** *p*_Bonf_ *< .*010).

### 3.2 Hyperscanning effect: true versus surrogate dyads

To determine whether the interpersonal neural synchrony observed during the Jenga task exceeded the level of coherence expected in the absence of direct interaction, we compared true and surrogate dyads separately across the 10 ROI pairings. Interpersonal neural synchrony was significantly higher in true than in surrogate dyads for all ROI pairings, with estimated differences in WTC ranging from *β* = 0.0030 to *β* = 0.0047 (*t* = 10.27–21.53, all Bonferroni-corrected *p*s *<* .001; see Table 2). Standardized effect sizes were relatively consistent across regional combinations, ranging from *d* = 0.35 to *d* = 0.55, indicating small-to-moderate differences in synchrony between true and surrogate dyads. Thus, the increased inter-brain coherence observed during the Jenga task was consistently present across the investigated regional combinations rather than being restricted to a single ROI pair.

### 3.3 Whole-network association between neural synchrony and Jenga performance

In the second stage of analysis, we examined whether behavioral performance during the Jenga task was associated with interpersonal neural synchrony across the fronto-temporal network. Interpersonal neural synchrony was entered as the dependent variable, while tower collapse and the number of valid turns completed during the task were included simultaneously as predictors. Neither behavioral measure showed a statistically significant association with interpersonal neural synchrony at the whole-network level. Tower collapse was associated with a small decrease in synchrony, but the effect did not reach statistical significance (*β* = *−*0.0017, *SE* = 0.0009, *t*(131.1) = *−*1.98, *p* = .050). The number of valid turns was also not significantly associated with synchrony (*β* = *−*0.00004, *SE* = 0.00008, *t*(128.7) = *−*0.48, *p > .*050).

Consistent with the whole-sample analysis, neither predictor reached statistical significance when the models were fitted separately within close-friend, romantic-partner, and mother–child dyads (all Bonferroni-corrected *p*s *> .*050).

### 3.4 ROI-specific associations between neural synchrony and performance

In the third stage of analysis, we examined whether behavioral performance was associated with interpersonal neural synchrony within specific ROI pairings. As shown in Table 3, tower collapse was significantly associated with reduced synchrony in two TPJ-related pairings after Bonferroni correction across all 20 tests. Specifically, dyads whose tower collapsed showed lower synchrony in the left TPJ–left TPJ pairing (*β* = *−*0.00458, *SE* = 0.00135, *t*(118.5) = *−*3.40, *p*_Bonf_ = .009) and in the left TPJ–right TPJ pairing (*β* = *−*0.00361, *SE* = 0.00112, *t*(124.6) = *−*3.21, *p*_Bonf_ = .017). No other ROI-specific association with tower collapse survived correction for multiple comparisons, and the number of valid turns was not significantly associated with interpersonal neural synchrony in any ROI pairing (all *p*_Bonf_ *> .*050).

To further characterize the ROI-specific effects, we examined the left TPJ–left TPJ and left TPJ–right TPJ pairings separately within each cohort (Table 4). For the left TPJ–left TPJ pairing, tower collapse was associated with lower interpersonal neural synchrony among close-friend dyads (*β* = *−*0.00580, *SE* = 0.00206, *t*(62.66) = *−*2.82, *p*_Bonf_ = .020) and romantic-partner dyads (*β* = *−*0.00716, *SE* = 0.00256, *t*(32.92) = *−*2.79, *p*_Bonf_ = .026), whereas no significant association was observed in mother–child dyads (*p*_Bonf_ *> .*050). For the left TPJ–right TPJ pairing, the association between tower collapse and interpersonal neural synchrony did not remain significant in any cohort after Bonferroni correction (all *p*_Bonf_ *> .*050). The number of valid turns was not significantly associated with synchrony in either ROI pairing within any cohort (all *p*_Bonf_ *> .*050). These follow-up analyses therefore suggest that the negative association between tower collapse and left TPJ–left TPJ synchrony was evident in the two adult cohorts but not in mother–child dyads.

**Table 4.** Cohort-specific associations between behavioral performance and interpersonal neural synchrony in the left TPJ–left TPJ and left TPJ–right TPJ pairings. Tower collapse (*fallen* vs. *not fallen*) and the number of valid turns were entered simultaneously as predictors of wavelet transform coherence (WTC). *P* values were Bonferroni-corrected across the three cohorts within each ROI pairing and behavioral predictor. Significant effects after correction are shown in bold (* *p*_Bonf_ *< .*050).

| ROI pairing | Cohort | Tower collapse |  |  |  | Valid turns |  |  |  |
| --- | --- | --- | --- | --- | --- | --- | --- | --- | --- |
| | | $\beta$ | SE | $t$ | $p_{\text{Bonf}}$ | $\beta$ | SE | $t$ | $p_{\text{Bonf}}$ |
| Left TPJ – Left TPJ | Close friends | -0.00580 | 0.00206 | -2.82 | **.020\*** | -0.00020 | 0.00019 | -1.09 | > .050 |
|  | Romantic partners | -0.00716 | 0.00256 | -2.79 | **.026\*** | -0.00003 | 0.00027 | -0.12 | > .050 |
|  | Mother–child | 0.00271 | 0.00270 | 1.01 | > .050 | 0.00007 | 0.00026 | 0.25 | > .050 |
| Left TPJ – Right TPJ | Close friends | -0.00447 | 0.00191 | -2.34 | > .050 | -0.00022 | 0.00017 | -1.26 | > .050 |
|  | Romantic partners | -0.00376 | 0.00179 | -2.10 | > .050 | 0.00005 | 0.00019 | 0.28 | > .050 |
|  | Mother–child | 0.00033 | 0.00230 | 0.15 | > .050 | 0.00025 | 0.00022 | 1.11 | > .050 |

## 4 Discussion

The present study investigated the relationship between interpersonal neural synchrony and cooperative performance during a naturalistic Jenga task performed by close friends, romantic partners, and mother–child dyads. Across all regional pairings, true interacting dyads exhibited greater inter-brain coherence than surrogate dyads, confirming that the cooperative interaction was accompanied by meaningful interpersonal neural synchrony. However, interpersonal neural synchrony across the broader fronto-temporal network was not significantly associated with either measure of behavioral performance. Instead, performance-related effects emerged at a more regionally specific level: tower collapse was associated with reduced synchrony in the left TPJ– left TPJ and left TPJ–right TPJ pairings, whereas the number of valid turns was unrelated to synchrony across all regional combinations. Follow-up analyses further showed that, for the left TPJ–left TPJ pairing, the negative association between tower collapse and synchrony was evident in close-friend and romantic-partner dyads but not in mother–child dyads. These cohort-specific analyses suggest that the relationship between TPJ synchrony and cooperative performance may vary across relational and developmental contexts.

The regional specificity of these effects is consistent with the proposed role of the TPJ in social coordination. The TPJ is a key component of the social brain network and has been consistently implicated in mentalizing, intention understanding, perspective taking, and the representation of others’ actions and goals (Carter & Huettel, 2013; Saxe & Kanwisher, 2003; Schurz, Radua, Aichhorn, Richlan, & Perner, 2014). These functions are particularly relevant during cooperative joint action, in which individuals must continuously monitor their partner’s behavior, anticipate the consequences of their actions, and update their own behavior accordingly. Such demands are especially salient in Jenga, where each move alters the shared physical environment and constrains the actions available to the partner on subsequent turns. Theoretical accounts further suggest that the TPJ may contribute to social interaction by integrating information unfolding over multiple temporal scales. In particular, Wolff and Dumas (2026) propose that intrinsic neural timescales within the TPJ enable the integration of relatively fast action-related information with slower processes involved in mentalizing and the representation of others’ intentions. From this perspective, inter-personal neural synchrony may reflect the temporal alignment of these integrative processes across interacting partners. During Jenga, such alignment could support the continuous integration of a partner’s actions, their consequences for the evolving state of the tower, and the adjustments required for subsequent moves. Greater TPJ synchrony may therefore index more effectively coordinated monitoring and adaptation during joint action, thereby reducing the likelihood of coordination failures such as tower collapse. This interpretation is consistent with previous hyperscanning studies linking temporo-parietal and prefrontal inter-brain synchrony to successful cooperation and joint problem solving (Czeszumski et al., 2022; Nguyen et al., 2020; Zhou, Cheng, Liu, & Li, 2022).

Importantly, the association with performance was specific to tower collapse rather than to the number of valid turns completed. These two measures may capture different aspects of cooperative performance. The number of valid turns primarily reflects the amount of successful task progression achieved within the fixed interaction period, whereas tower collapse represents a more salient failure to maintain the shared physical structure of the game. Avoiding collapse requires participants not only to perform their individual actions successfully, but also to continuously take into account the consequences of previous moves and the changing state of the tower. Reduced TPJ synchrony in dyads whose tower collapsed may therefore reflect less effective alignment of the monitoring and anticipatory processes required to maintain coordinated action toward a shared external goal. This distinction may also help explain why previous studies have reported mixed and heterogeneous associations between interpersonal synchrony and joint performance (Gordon & Bartsch, 2026). Measures labeled as “performance” are not necessarily functionally equivalent: some capture overall task output or speed, whereas others index coordination failures, accuracy, successful completion, or the ability to maintain a shared goal over time. Interpersonal neural synchrony may be more closely related to some of these components than to others, depending on the cognitive and interactional demands of the task. In the present study, synchrony was associated with a discrete coordination failure, but not with the cumulative number of successful actions. Thus, part of the variability observed across studies may arise not only from differences in tasks and dyadic composition, but also from differences in how cooperative performance is operationalized.

The cohort-specific follow-up analyses provide further insight into the context dependence of the synchrony–performance relationship. In the left TPJ–left TPJ pairing, tower collapse was associated with lower interpersonal neural synchrony among both close friends and romantic partners, whereas no corresponding association was observed among mother–child dyads. Although these analyses do not constitute a formal test of differences between cohorts, the pattern is consistent with the broader proposal that the behavioral relevance of interpersonal neural synchrony may depend on who is interacting (Carollo, Bizzego, et al., 2025). Social relationships differ in shared history, familiarity, interpersonal expectations, and established patterns of coordination, all of which can influence how partners anticipate and respond to one another during joint action. Previous hyperscanning findings provide support for this possibility. Romantic relationships, in particular, have been associated with both enhanced cooperative performance and distinct patterns of inter-brain synchrony. For example, romantic partners have been shown to display better cooperative behavior and stronger interpersonal neural synchrony than friends or unfamiliar partners, with synchrony covarying with cooperative performance (Pan et al., 2017). Similarly, during collaborative creative problem solving, romantic partners outperformed stranger dyads in measures of cooperation and task performance and exhibited greater interpersonal synchrony in frontopolar and temporo-parietal regions. Within romantic partners, stronger synchrony was also associated with better cooperative performance (Duan et al., 2022). These examples suggest that interpersonal neural synchrony cannot be interpreted independently of the relational context in which coordination occurs. A history of repeated interaction and shared experience may shape how efficiently partners predict, interpret, and adapt to one another, potentially altering the relationship between inter-brain coupling and successful joint behavior.

In the present study, however, similar synchrony–performance patterns were observed in both close friends and romantic partners, whereas the association was absent in mother–child dyads. This suggests that the relevant distinction may not be romantic involvement itself, but a broader combination of relational and developmental characteristics that distinguishes adult–adult from adult–child interaction. Parent– child interactions are embedded within an asymmetric developmental relationship in which partners differ substantially in cognitive, motor, and regulatory abilities and in which the adult frequently takes on a scaffolding role. Moreover, shared activities in parent–child dyads can serve functions beyond objective task completion, including mutual engagement, socio-emotional connection, learning, and regulation (Yogman et al., 2018). Interpersonal neural synchrony in such interactions may therefore track aspects of responsiveness, engagement, or affective attunement that are only partially reflected in objective game outcomes. This interpretation is compatible with evidence that parent–child neural synchrony varies with features of interaction quality and interpersonal engagement during cooperative activities (Nguyen et al., 2020; Nguyen, Schleihauf, Kayhan, et al., 2021; Nguyen, Schleihauf, Kungl, et al., 2021).

Developmental differences in task demands may provide a complementary explanation. Successful Jenga performance relies not only on interpersonal coordination but also on fine motor control, visuomotor integration, and the ability to anticipate the physical consequences of object manipulation. These abilities are still developing during early childhood (Adolph & Franchak, 2017), and motor performance remains comparatively variable during the preschool years (Sutapa, Pratama, Rosly, Ali, & Karakauki, 2021). Tower collapse may therefore not constitute an equivalent measure of cooperative success across adult and mother–child dyads. In adult dyads, a collapse may be more closely related to failures in planning or joint coordination, whereas in mother–child dyads it may also reflect developmental limitations in motor precision that are partly independent of interpersonal coordination. Moreover, mother–child cooperation is inherently more asymmetric than cooperation between adult partners. Mothers may adapt their behavior to the child’s abilities, provide greater scaffolding, and place relatively greater emphasis on maintaining the child’s engagement and participation than on maximizing task performance. Consequently, the interactional meaning of a tower collapse may differ across dyad types even when the observable event is the same. Under this interpretation, the absence of a synchrony–collapse association in mother–child dyads would not imply weaker cooperation, but rather that both the behavioral demands of the task and the organization of the interaction differ from those characterizing adult–adult dyads.

Overall, the present findings highlight the importance of considering neural synchrony at an appropriate spatial, behavioral, and relational level. Although interacting dyads showed greater synchrony than surrogate dyads throughout the measured network, only specific TPJ pairings were associated with task outcome. Interpersonal neural synchrony may therefore reflect multiple processes during social interaction, only some of which are directly related to behavioral success. The absence of a whole-network association, together with the regionally and behaviorally specific TPJ findings, is consistent with theoretical accounts that conceptualize interpersonal synchrony as flexible and context dependent rather than uniformly adaptive (Gordon et al., 2025). From this perspective, the behavioral relevance of interpersonal neural synchrony depends on the neural systems involved, the demands of the interaction, the outcome used to define successful coordination, and the characteristics of the interacting partners (Carollo, Bizzego, et al., 2026).

### 4.1 Limitations

Some limitations should be considered when interpreting the present findings.

First, the observational nature of the study precludes causal conclusions regarding the relationship between interpersonal neural synchrony and cooperative performance (Novembre & Iannetti, 2021). Although greater left TPJ synchrony was associated with a lower likelihood of tower collapse in the adult cohorts, the direction of this relationship cannot be established. Neural alignment may contribute to successful coordination, successful coordination may promote neural alignment, or both may reflect broader interactional processes.

Second, the Jenga task may not have represented an equivalent challenge across participant groups. Successful performance depends not only on interpersonal coordination but also on fine motor control, visuomotor integration, and manual dexterity. Given that children in the mother–child cohort were approximately five years old, developmental differences in these abilities may have contributed to tower collapses independently of interpersonal coordination. Accordingly, caution is warranted when directly comparing the behavioral significance of tower collapse across adult and mother–child dyads.

Finally, cooperation was assessed within a single naturalistic task using two behavioral indices. Although tower collapse and the number of valid turns provide objective measures of game performance, they capture different and only partially overlapping aspects of cooperative success. The present findings should therefore be interpreted in relation to the specific behavioral and interactional demands of the Jenga task.

## 5 Conclusion

The present study investigated the relationship between interpersonal neural synchrony and cooperative performance during a naturalistic Jenga task across different social relationships (Carollo, Bizzego, et al., 2025). Consistent with previous hyper-scanning research, true interacting dyads exhibited greater inter-brain synchrony than surrogate dyads across all examined regional pairings, indicating that real-time interaction was accompanied by enhanced neural synchrony.

Importantly, the association between neural synchrony and cooperative performance was regionally specific. Tower collapse was associated with reduced synchrony involving the left TPJ, whereas the number of valid turns was unrelated to neural synchrony. Follow-up analyses further showed that lower left TPJ–left TPJ synchrony was associated with tower collapse among close friends and romantic partners, while no corresponding association emerged in mother–child dyads.

Together, these findings suggest that the behavioral relevance of interpersonal neural synchrony is context-dependent rather than uniform across the brain, performance measures, and interacting partners (Gordon & Bartsch, 2026; Gordon et al., 2025). In adult dyads, TPJ synchrony may reflect the coordinated social-cognitive processes involved in monitoring a partner and adapting behavior toward a shared external goal. In mother–child dyads, by contrast, the relationship between neural coupling and objective game performance may be shaped by a broader combination of relational, developmental, and task-specific factors.

Overall, the present results highlight the importance of considering what partners are doing, how successful coordination is operationalized, and who is interacting when interpreting interpersonal neural synchrony. Rather than serving as a general marker of successful cooperation, interpersonal neural synchrony may be most informative when considered in relation to specific neural systems and the social and behavioral context in which joint action unfolds.

## Authors contribution

Conceptualization: AC, AB, RC, DD, SH, GE; Data Curation: AC, RC, AB; Formal analysis: AC, RC, AB; Investigation: AC, RC; Methodology: AC, AB; Supervision: SH, GE; Writing–original draft preparation: AC, RC; Writing–review and editing: AC, RC, AB, DD, SH, GE. All authors have read and agreed to the published version of the manuscript.

