## Supplementary Materials for "Left Temporoparietal Neural Synchrony Tracks Cooperative Performance Across Adult Dyads"

### S1. Hyperscanning effect: true versus surrogate dyads

| ROI pairing | $\beta$ | SE | $t$ | df | $p_{\mathrm{Bonf}}$ | $d$ [95% CI] |
|---|---|---|---|---|---|---|
| Left IFG – Left IFG | 0.00926 | 0.00068 | 13.71 | 3934.7 | < .001*** | 0.43 [0.37, 0.49] |
| Left IFG – Right IFG | 0.01183 | 0.00051 | 23.32 | 7880.1 | < .001*** | 0.52 [0.48, 0.57] |
| Left IFG – Left TPJ | 0.01082 | 0.00052 | 20.92 | 7776.0 | < .001*** | 0.47 [0.43, 0.52] |
| Left IFG – Right TPJ | 0.00823 | 0.00064 | 12.85 | 4583.5 | < .001*** | 0.38 [0.32, 0.43] |
| Right IFG – Right IFG | 0.01381 | 0.00073 | 18.90 | 3832.8 | < .001*** | 0.61 [0.54, 0.67] |
| Right IFG – Left TPJ | 0.01422 | 0.00052 | 27.27 | 7679.2 | < .001*** | 0.62 [0.58, 0.67] |
| Right IFG – Right TPJ | 0.01130 | 0.00065 | 17.27 | 4516.5 | < .001*** | 0.51 [0.45, 0.57] |
| Left TPJ – Left TPJ | 0.01328 | 0.00068 | 19.48 | 3730.2 | < .001*** | 0.64 [0.57, 0.70] |
| Left TPJ – Right TPJ | 0.01193 | 0.00067 | 17.71 | 4442.7 | < .001*** | 0.53 [0.47, 0.59] |
| Right TPJ – Right TPJ | 0.00875 | 0.00111 | 7.86 | 1349.7 | < .001*** | 0.42 [0.22, 0.61] |

Table S1: Comparison of interpersonal neural synchrony between true and surrogate dyads across ROI pairings during the cooperative Jenga task, based on deoxygenated hemoglobin (HbR) signals. Positive coefficients indicate greater wavelet transform coherence (WTC) in true compared with surrogate dyads. $P$ values were Bonferroni-corrected across the 10 ROI comparisons. (*** $p_{\mathrm{Bonf}} < .001$).

### S2. Whole-network association between neural synchrony and Jenga performance

| | Tower collapse | | | | | Valid turns | | | | |
|---|---|---|---|---|---|---|---|---|---|---|
| **Sample** | $\beta$ | SE | $t$ | df | $p$ | $\beta$ | SE | $t$ | df | $p$ |
| Overall sample | -0.00032 | 0.00219 | -0.15 | 129.3 | > .050 | 0.00011 | 0.00022 | 0.50 | 128.8 | > .050 |
| Close friends | -0.00250 | 0.00351 | -0.71 | 64.4 | > .050 | 0.00024 | 0.00031 | 0.76 | 64.0 | > .050 |
| Romantic partners | -0.00248 | 0.00461 | -0.54 | 34.0 | > .050 | -0.00024 | 0.00054 | -0.44 | 34.0 | > .050 |
| Mother–child | 0.00781 | 0.00314 | 2.49 | 27.2 | > .050 | 0.00047 | 0.00031 | 1.55 | 26.8 | > .050 |

Table S2: Associations between behavioral performance and interpersonal neural synchrony at the whole-network level based on deoxygenated hemoglobin (HbR) signals. Tower collapse (*fallen* vs. *not fallen*) and the number of valid turns were entered simultaneously as predictors of wavelet transform coherence (WTC). For the overall sample, uncorrected $p$ values are reported. For cohort-specific analyses, $p$ values were Bonferroni-corrected separately for each behavioral predictor across the three cohorts. No cohort-specific effects remained significant after correction.

### S3. ROI-specific associations between neural synchrony and performance

| ROI pairing | Tower collapse | | | | Valid turns | | | |
|---|---|---|---|---|---|---|---|---|
| | $\beta$ | SE | $t$ | $p_{\mathrm{Bonf}}$ | $\beta$ | SE | $t$ | $p_{\mathrm{Bonf}}$ |
| Left IFG – Left IFG | -0.00305 | 0.00249 | -1.22 | $> .050$ | 0.00056 | 0.00026 | 2.14 | $> .050$ |
| Left IFG – Right IFG | 0.00114 | 0.00247 | 0.46 | $> .050$ | 0.00026 | 0.00025 | 1.07 | $> .050$ |
| Left IFG – Left TPJ | 0.00009 | 0.00266 | 0.03 | $> .050$ | 0.00013 | 0.00026 | 0.50 | $> .050$ |
| Left IFG – Right TPJ | -0.00635 | 0.00260 | -2.44 | $> .050$ | 0.00010 | 0.00027 | 0.38 | $> .050$ |
| Right IFG – Right IFG | -0.00080 | 0.00341 | -0.23 | $> .050$ | 0.00008 | 0.00035 | 0.24 | $> .050$ |
| Right IFG – Left TPJ | 0.00044 | 0.00287 | 0.15 | $> .050$ | -0.00017 | 0.00028 | -0.59 | $> .050$ |
| Right IFG – Right TPJ | -0.00243 | 0.00282 | -0.86 | $> .050$ | -0.00019 | 0.00030 | -0.62 | $> .050$ |
| Left TPJ – Left TPJ | 0.00102 | 0.00325 | 0.32 | $> .050$ | -0.00009 | 0.00032 | -0.29 | $> .050$ |
| Left TPJ – Right TPJ | -0.00026 | 0.00302 | -0.09 | $> .050$ | 0.00022 | 0.00031 | 0.71 | $> .050$ |
| Right TPJ – Right TPJ | -0.00396 | 0.00346 | -1.15 | $> .050$ | 0.00011 | 0.00038 | 0.28 | $> .050$ |

Table S3: Associations between behavioral performance and interpersonal neural synchrony across ROI pairings based on deoxygenated hemoglobin (HbR) signals. Tower collapse (*fallen* vs. *not fallen*) and the number of valid turns were entered simultaneously as predictors of wavelet transform coherence (WTC). $P$ values were Bonferroni-corrected separately for each behavioral predictor across the 10 ROI pairings. No effects remained significant after correction.
